# Signatures for Viral Infection and Inflammation in the Proximal Olfactory System in Familial Alzheimer’s Disease

**DOI:** 10.1101/2022.07.19.500641

**Authors:** Andrew N. Bubak, Laetitia Merle, Christy S. Niemeyer, B. Dnate’ Baxter, Arianna Gentile Polese, Vijay Ramakrishnan, Johana Gomez, Lucia Madrigal, Andres Villegas-Lanau, Francisco Lopera, Wendy Macklin, Seth Frietze, Maria A. Nagel, Diego Restrepo

## Abstract

**Objective:** Alzheimer’s disease (AD) is characterized by loss of smell and olfactory system pathology that precedes the diagnosis of dementia. Understanding these early processes can potentially identify diagnostic and therapeutic targets to slow AD progression. Here we analyzed differential gene and protein expression in the olfactory bulb (OB) and olfactory tract (OT) of familial AD (FAD) individuals carrying the autosomal dominant presenilin 1 E280A paisa mutation and age-matched controls.

**Methods:** Formalin-fixed, paraffin-embedded sections containing both the OB and OT from 6 FAD individuals and 6 age-matched controls were obtained. Tissue morphology and composition were characterized by immunohistochemistry using antibodies against the myelin marker proteolipid protein (PLP), amyloid-beta (Aβ), and microglia/macrophage markers Iba1 and CD68, respectively. OB and OT were analyzed separately by targeted RNA sequencing of the whole human transcriptome (BioSpyder TempO-Seq); ingenuity pathway analysis and R-computational program were used to identify differentially expressed genes and pathways between groups. The nanoString spatial proteomics assay for 88 proteins, including markers for AD and immune responses, was used to complement gene expression findings.

**Results:** Compared to control OT, FAD OT had significantly increased immunostaining for Aβ and CD68 in the high and low myelinated regions, as well as increased immunostaining for Iba1 in the high myelinated region only; both control and FAD OT samples had similar total area of high and low myelinated regions. In FAD samples, RNA sequencing showed a transcription profile consistent with: (1) viral infection in the OB; (2) inflammation in the OT that carries information via entorhinal cortex from the OB to hippocampus, a brain region essential for learning and memory; and (3) decreased oligodendrocyte deconvolved transcripts, indicating dysregulation of myelination. Interestingly, spatial proteomic analysis confirmed altered myelination in the OT of FAD individuals, implying dysfunction of communication between the OB and hippocampus.

**Conclusions:** These findings raise the possibility that viral infection and associated inflammation and dysregulation of myelination of the olfactory system may disrupt downstream hippocampal function, contributing to acceleration of FAD progression.

## Introduction

A critical barrier in treating Alzheimer’s disease (AD) is the years- to decades-long lag from disease onset to the diagnosis of dementia, when reversal of brain pathology and recovery of neurons may, at best, slow cognitive decline. Thus, it is essential to identify contributory pathological processes in early disease to prevent progression to dementia, disability, and death. An early process in AD, prior to clinical dementia, is a deficit in the sense of smell^1-3^ accompanied by amyloid-beta (Aβ) deposition in the olfactory bulb (OB)^4, 5^. The OB receives olfactory input from olfactory sensory neurons in the olfactory epithelium (OE)^3, 6^ then transmits olfactory information via the olfactory tract (OT) to the entorhinal cortex and hippocampus, brain regions essential for learning and memory that are affected in AD. Dissecting mechanisms involved in olfactory dysfunction can identify potential diagnostic and therapeutic targets in AD. However, little is known about changes in the cellular processes of the olfactory system, including the OB and OT.

Because sporadic AD is quite heterogenous due to variable contributions of host and environmental factors, we focused our initial studies on olfactory tissues from a well-characterized familial AD (FAD) cohort from Colombia. These FAD individuals have the presenilin 1 E280A mutation and develop mild cognitive impairment (MCI) and dementia at the respective median ages of 44 (95% CI, 43–45) and 49 (95% CI, 49–50) years old^7^. Formalin-fixed paraffin embedded (FFPE) slides containing olfactory tissue collected from this FAD cohort post-mortem, as well as control tissue, were examined for tissue morphology and composition. Adjacent slides were analyzed by targeted RNA sequencing of the OB and OT (TempO-Seq, BioSpyder Technologies, Inc., Carlsbad, CA)^8^ and by spatial proteomics (nanoString Technologies, Seattle, WA)^9^; bioinformatics analyses were used to identify significantly enriched gene expression pathways.

## Methods

### Standard Protocol Approvals, Registrations, Patient Consents

Tissue from individuals in Colombia with the presenilin 1 PSEN1 E280A mutation ^10^ and age-matched controls were collected for this study. Approval for post-mortem studies was granted by the institutional review board committee at the University of Antioquia (Comité de Bioetica de la Sede de Investigación Universitaria, Universidad de Antioquia, Colombia) on June 26th, 2019 (approval number 19010-858). Participants provided written informed consent before participating, agreeing to donate tissue for research at time of death. A total of 6 members from Colombian families with the *PSEN1* E280A mutation and 6 age-matched controls were available for analysis in the study. Demographic data and associated studies completed herein are detailed in Table 1. The average age (range, years [y]) of the control and FAD groups was 69 y (56-84 y) and 62 y (53-86 y), respectively. The control group contained 5 males and 1 female; the FAD group contained 2 males and 4 females. None of the individuals in either control of FAD groups died of viral infection that may affect interpretation of results.

**Table 1.** Patient Information.

| Subject ID | Group | Age | Sex | Cause of Death | OT surface analyzed in Vectra (mm<sup>2</sup>) | Temp-O-SEQ OT | Temp-O-SEQ OB | Nanostring OT | Nanostring OB |
| --- | --- | --- | --- | --- | --- | --- | --- | --- | --- |
| 337 | Control | 84 | M | Bacterial pneumonia | 5.1 | X | X | X | X |
| 351 | Control | 62 | M | Heart failure | 7.0 |  | X |  |  |
| 354 | Control | 65 | M | Decompensated chronic obstructive pulmonary disease | 3.3 |  | X |  |  |
| 360 | Control | 72 | M | Acute myocardial infarction | 2.1 | X |  |  |  |
| 364 | Control | 73 | M | Gastric cancer | 3.9 | X |  |  |  |
| 380 | Control | 56 | F | Colon cancer | 4.1 | X | X | X | X |
| 359 | FAD (E280A) | 56 | F | Bacterial pneumonia | 3.6 | X | X | X | X |
| 346 | FAD (E280A) | 53 | M | Aspiration pneumonia | 5.2 | X | X |  |  |
| 355 | FAD (Late Onset) | 86 | F | Acute renal failure | 3.0 | X | X |  |  |
| 335 | FAD (E280A) | 59 | M | Bacterial pneumonia | 2.7 | X | X |  |  |
| 358 | FAD (E280A) | 59 | F | Aspiration pneumonia | 1.1 | X | X | X | X |
| 382 | FAD (E280A) | 57 | F | Metabolic imbalance | 3.8 | X | X |  |  |

The E280A carriers have a previously documented median expected age at onset of MCI at 44 years of age (95% confidence interval [CI] = 43–45 years) and dementia due to AD at 49 years of age (95% CI = 49–50 years) ^7^. Eligible individuals were screened for neurological and psychiatric disorders prior to death, and they did not die of viral pneumonia. At time of death, all 6 FAD individuals had a clinical diagnosis of dementia. Clinical data were stored in a secure database at the Neuroscience Group of Antioquia in Medellín, Colombia.

### Sample Collection

Brain samples were collected post-mortem; tissue containing the OB and OT were fixed in formalin then paraffin-embedded. OB/OT tissue blocks were cut sagittally at 7 μm thickness and placed on slides for hematoxylin and eosin staining (H&E; data not shown) and immunohistochemical (IHC) staining.

### Multispectral Immunohistochemistry of the Olfactory Bulb and Olfactory Tract

Through our collaboration with the Human Immune Monitoring Shared Resource (HIMSR) at the University of Colorado School of Medicine, we performed multispectral imaging using the Akoya Vectra Polaris instrument. This instrumentation allows for phenotyping, quantification, and spatial relationship analysis of tissue infiltrate in formalin-fixed paraffin-imbedded biopsy sections^11^.

FFPE tissue sections containing OB and OT were stained consecutively with specific primary antibodies according to standard protocols provided by Akoya and performed routinely by the HIMSR. Briefly, the slides were deparaffinized, heat treated in antigen retrieval buffer, blocked, and incubated with primary antibodies against Aβ, Iba1, GFAP, Cleaved Caspase 3, p-Tau, DCX, CD68, and PLP (Supplemental Table 1), followed by horseradish peroxidase (HRP)-conjugated secondary antibody polymer, and HRP-reactive OPAL fluorescent reagents that use TSA chemistry to deposit dyes on the tissue immediately surrounding each HRP molecule. To prevent further deposition of fluorescent dyes in subsequent staining steps, the slides were stripped in between each stain with heat treatment in antigen retrieval buffer. Slides were finally counterstained with DAPI. Whole slide scans were collected using the 10x objective and multispectral images were collected using the 20x objective with a 0.5 micron resolution. The 9-color images were analyzed with inForm software V2.5 to unmix adjacent fluorochromes, subtract autofluorescence, segment the tissue, compare the frequency and location of cells, segment cellular membrane, cytoplasm, and nuclear regions, score each cellular compartment, and phenotype infiltrating immune cells according to morphology and cell marker expression.

We used custom scripts in Fiji to quantify Aβ, Iba1, CD68 and PLP on the 20x objective images; GFAP, Cleaved Caspase 3, p-Tau, and DCX antibodies did not yield reliable staining and were not further analyzed. ROIs for high and low myelinated areas were manually determined using PLP stain by two researchers. To quantify Aβ, Iba1 and CD68-positive areas, we defined the positive area as one that exceeds *n* SDs of the mean image intensity, with *n* = 2 for Aβ and Iba1 and *n* = 4 for CD68. The positive surface was expressed as a percentage of the ROI surface per individual. Masks for each stain were created from the thresholded images, and mask combinations were used to determine the double and triple-positive areas.

### FFPE TempO-Seq

For whole human transcriptome analysis, FFPE slides containing OB or OT were assayed using TempO-Seq–targeted RNA sequencing plates, reagents, protocols, and software (BioSpyder Technologies)^8^. TempO-Seq exclusively detects human transcripts; viral transcripts were not assayed. H&E stained tissue sections were used to distinguish the OB from OT. Of the 6 controls, only 4 had adequate samples for this analysis whereas all 6 FAD samples contained OB and OT for this analysis. The OB and OT from adjacent, non-stained slides were separately scraped (~5-10 mm^2^) and placed into PCR tubes containing 1X lysis buffer. Using a thermocycler, samples containing OB or OT were lysed. Coded adjacent primer pairs for each specific human transcript were annealed to sample RNA; primer pairs for each transcript were ligated and then amplified per manufacturer’s instructions and as previously described ^12^. If a transcript is present, the adjacent 25-nucleotide primer pairs anneal to their specific target, ligate together, and produce a 50-nucleotide transcript-specific, coded amplicon.

Amplified PCR products were pooled into a single library, concentrated using a PCR cleanup kit (Macherey-Nagel, Düren, Germany), and run on the Illumina NextSeq 500 sequencing platform (Illumina Inc., San Diego, CA). Mapped reads were generated by TempO-SeqR for the alignment of demultiplexed FASTQ files from the sequencer to the ligated detector oligomer gene sequences using Bowtie, allowing for up to 2 mismatches in the 50-nucleotide target sequence^8^. Counts were assessed using SARTools^13^. Within this R package, edgeR is used for normalization and quality control of count data^14^. Differential expression between groups was assessed by the TempO-SeqR software, which used the DESeq2 method for differential analysis of count data^15^. A significantly differentially expressed gene is defined as having an adjusted p value < 0.05 with no fold-change threshold.

Pathway enrichment analysis was performed using gene sets and pathways defined in the Ingenuity Pathway Analysis software (Qiagen, Germantown, MD) and the ClusterProfiler package^16, 17^ in R with default parameters. ClusterProfiler supports enrichment analysis of Gene Ontology and Kyoto Encyclopedia of Genes and Genomes databases with Gene Set Enrichment Analysis (GSEA) to identify biological themes of a collection of genes. These functional enrichment analyses use computational approaches to identify groups of experimentally observed human genes that are overrepresented or depleted in a curated disease or biological function-specific gene set. Additional figures were created using Prism 9 (GraphPad Software, San Diego, CA).

### nanoString Proteomics

We used nanoString proteomics for 88 proteins (Supplemental Table 2 with four fluorescent markers to discriminate tissue regions (Supplemental Table 3) using methods detailed by Merritt and co-workers^9^. Briefly, all assays were performed on 7-μm FFPE sections mounted onto charged slides; these slides were adjacent to those used for TempO-Seq analysis. Deparaffinization and rehydration of tissue was performed by incubating slides in three washes of CitriSolv (Decon Labs, 1601) for 5 minutes each, two washes of 100% ethanol for 10 minutes each, two washes of 95% ethanol for 10 minutes each and two washes of distilled water for 5 minutes each. For antigen retrieval, slides were then placed in a plastic Coplin jar containing 1× Citrate Buffer pH 6.0 (Sigma, C9999) and covered with a lid. The Coplin jar was placed into a pressure cooker (BioSB, BSB7008) that was run at high pressure and temperature for 15 minutes. The Coplin jar was removed from the pressure cooker and cooled at room temperature for 25 minutes. Slides were washed with five changes of 1× TBS-T (Cell Signaling Technology, 9997) for 2 minutes each. Excess TBS-T was removed from the slide, and a hydrophobic barrier was drawn around each tissue section with a hydrophobic pen (Vector Laboratories, H-4000). Slides were then incubated with blocking buffer (1× TBS-T, 5% goat serum (Sigma-Aldrich, G9023-5ML), 0.1 mg ml–1 salmon sperm DNA (Sigma-Aldrich, D7656) and 10 mg ml–1 dextran sulfate (Sigma-Aldrich, 67578-5G) for 1 hour. Slides were washed with three changes of 1× TBS-T for 2 minutes each. Primary antibodies (Supplemental Tables 2 and 3) were diluted in Buffer W (GeoMx Protein Slide Prep Kit Item 121300312). Tissue sections were then covered with diluted primary antibody solution (see above for morphology marker information). Slides were incubated at 4°C in a humidity chamber overnight. Primary antibody was aspirated from slides, and slides were washed with three changes of 1× TBS-T for 10 minutes each. Antibodies were postfixed with 4% paraformaldehyde for 30 minutes at room temperature and then wasted twice in TBS-T. DNA was counterstained with 500 nM SYTO13 (Thermo Fisher, S7575) in 1× TBS-T or TBS for 15 minutes. Excess DNA counterstain was removed with five changes of TBS-T, and slides were the processed on the GeoMx instrument as described in Merritt et al.^9^.

## Statistical Analysis

Matlab or Excel were used to perform t tests or ANOVAs, followed by post-hoc Fisher’s least significant difference procedure.

## Data Availability

All data used in the analyses of this study are available within the manuscript and its supplemental information files. Raw gene counts generated from the TempO-Seq analysis are deposited in the NCBI Gene Expression Omnibus database. Requests for further data sharing including individual patient sample information will be reviewed by the corresponding author and appropriate regulatory bodies with respect to maintaining de-identified status.

## Results

### Immunohistochemical analysis shows Aβ and microglia/macrophages in FAD olfactory tissue

Prior to transcriptome analysis, we analyzed the tissue for morphology and composition. Figure 1 A-C shows immunohistochemical analysis of the human OT from FAD subjects and age-matched controls (Table 1). Staining for the myelin marker proteolipid protein (PLP)^18^ showed that the OT included high and low myelination areas (Figure 1A). We did not find a difference between FAD and controls in the area occupied by the high/low myelinated regions in the OT (Figure 1A). As expected, we found increased immunostaining for Aβ in the OT of FAD subjects compared to controls (Figure 1B). In addition, as expected for neuroinflammation in AD, and consistent with prior studies in spontaneous AD^19^, we found increased immunostaining for the microglia/macrophage markers Iba1 and CD68 in FAD OT compared to control (Figure 1C). Finally, we found increased association between CD68 and Iba1 and between Aβ and microglia whose association has been proposed to constitute a barrier with profound impact on Aβ plaque composition and toxicity^20^ (Figure 1D-E).

**Figure 1.**
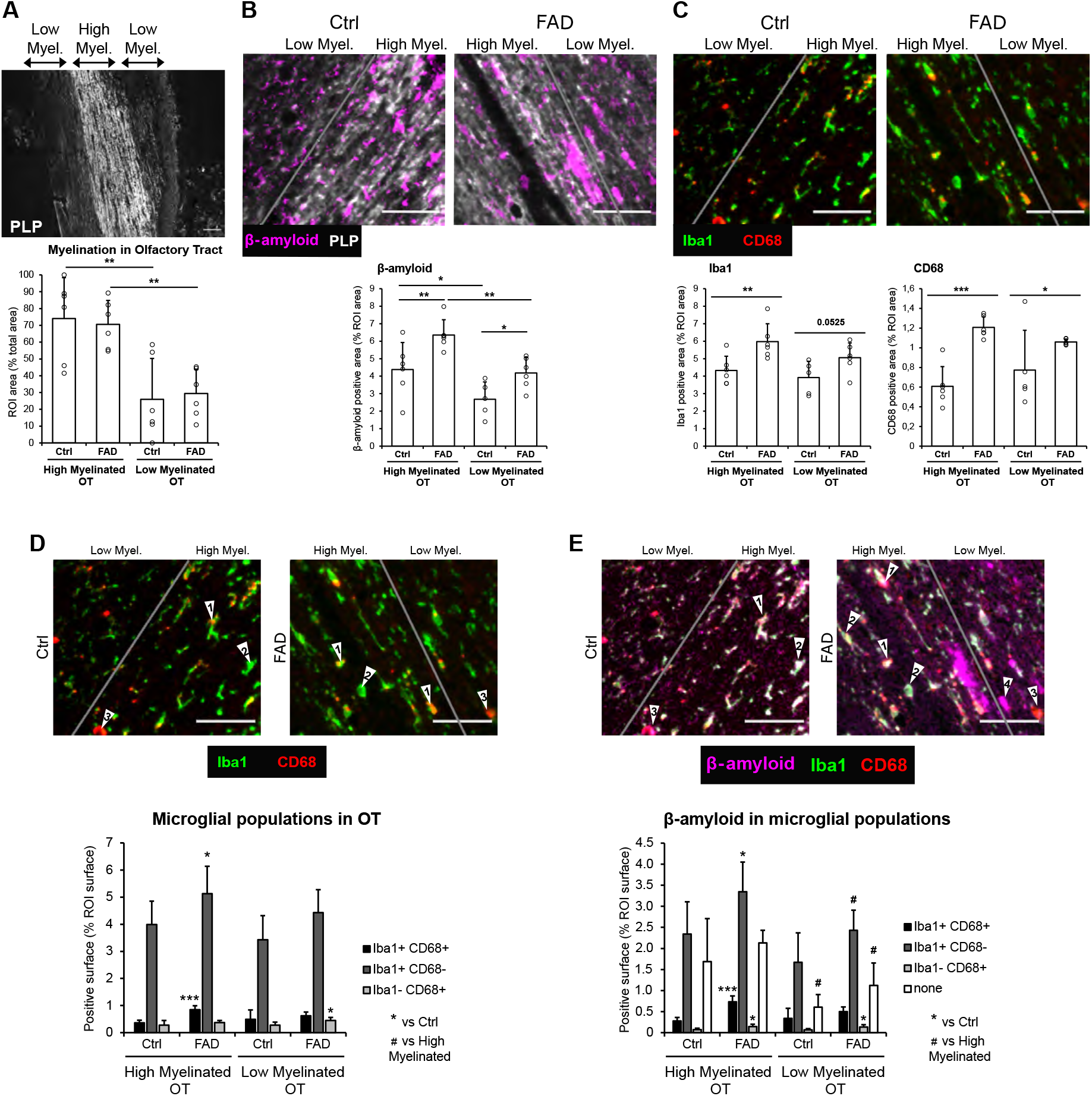
Immunohistochemistry of the Human OT and TempO-Seq Transcriptome Analysis of the OB and OT (A-C) IHC of the human OT for 6 control and 6 FAD subjects. (A) PLP stain was used to delimitate the high and low myelinated regions in OT. The bar graph represents the mean surface of high and low myelinated regions per individual, expressed as a percentage of the total area analyzed. An ANOVA yields no significant difference for the genotype F=0.1, 1 d.f., *p*>0.05 and a significant difference between high and low myelination areas F=26.84, 1 d.f., *p*<0.001. (B) Representative images and quantification of Aβ in the OT from 6 Ctrl and 6 FAD patients. Aβ positive surface is increased in FAD patients compared to control, in both the high and low myelinated OT. Within the FAD group, the accumulation of Aβ is higher in the high myelinated than the low myelinated OT. ANOVA yields a significant difference for the genotype F=13.94, 1 d.f., *p*<0.01 and a significant difference between high and low myelination areas F=17.22, 1 d.f., *p*<0.001. (C) Quantification of microglia/macrophage markers Iba1 and CD68, showing increased neuroinflammation in the high myelinated OT. ANOVA for Iba1 and CD68 yields a significant difference for genotype (Iba1, F=13.54, 1 d.f., *p*<0.01, CD68, F=23.24, 1 d.f., *p*<0.001). (D and E) Quantification of overlap in expression of Aβ, Iba1 and CD68. (D) Quantification of the 3 different microglial populations: Iba1+CD68+ (arrows “1”), Iba1+CD68- (arrows “2”), Iba1-CD68+ (arrows “3”). An ANOVA yields a significant difference for the genotype for Iba1+CD68+ (F=14.1, 1 d.f., p<0.01), Iba1+CD68- (F=8, 1 d.f., p<0.05) and Iba1-CD68+ (F=6.65, 1 d.f., p<0.05). FAD patients exhibit an increase in Iba1+CD68+ surface and Iba1+CD68-surface in the high myelinated OT, and an increase in Iba1-CD68+ surface low myelinated OT, compared to Ctrl. (E) Quantification of Aβ surface that is overlapping with Iba1+CD68+ cells. An ANOVA yields a significant difference for the genotype for Aβ surface that is overlapping with Iba1+CD68+ cells (F=24.14, 1 d.f., p<0.001), Iba1+CD68-cells (F=9.93, 1 d.f., p<0.01) and Iba1-CD68+ cells (F=12.28, 1 d.f., p<0.01). An ANOVA yields a significant difference between high and low myelination areas for Aβ surface that is overlapping with Iba1+CD68-cells (F=8, 1 d.f., p<0.05) and Aβ free from microglial coverage (F=15.93, 1 d.f., p<0.001). Aβ surface that is overlapping with all three microglial populations is increased in the high myelin OT of FAD patients compared to Ctrl. Aβ surface that is overlapping with Iba1-CD68+ cells is increased in the low myelin OT. Aβ free from microglial coverage (arrows “4”) is decreased in the low myelin OT compared to the high myelin OT in both Ctrl and FAD. Scale bar: 50 µm.

### Targeted RNA sequencing shows enrichment of pathways involved in virus infection in OB and of inflammation in OT

We then performed TempO-seq targeted RNA sequencing analysis of the OB and OT from 6 FAD individuals who did not die of virus infection and 4 age-matched controls (Table 1). Compared to controls, we found 2,775 differentially expressed genes (DEGs) in FAD OB and 1,319 significantly DEGs in FAD OT. Volcano plots in Figures 2Ai (OB) and 2Aii (OT) show that the majority of DEGs displayed increased expression in FADs (also see Supplemental Tables 4 and 5). Human gene-set enrichment analyses in the FAD OB showed multiple upregulated transcriptional pathways associated with viral infection, including “Viral infection”, “Infection of cells”, and “Replication of virus” (Figure 2Aiii), whereas the OT of FAD individuals represented an upregulation of immune response signatures that included “IL-6 signaling”, “IL-8 signaling”, “Natural killer cell signaling”, “Jak/Stat signaling”, “Neuroinflammation signaling”, and “Production of nitric oxide and reactive oxygen species in macrophages,” as well as the expected “Amyloid processing” pathway (Figure 2Aiv). Observed genes in each pathway are shown in Supplemental Table 6. When we deconvolved the DEGs to estimate glial surrogate proportions, we found that microglia were higher in FAD OT samples (Figure 2C), consistent with the immunohistochemistry results for Iba1 and CD68 (Figure 1C). Interestingly, in the deconvolution, we found a decreased surrogate proportion for oligodendrocytes in OT in FAD (Figure 2C). No differences in glial surrogate proportions were observed in FAD OB samples compared to controls (not shown).

**Figure 2.**
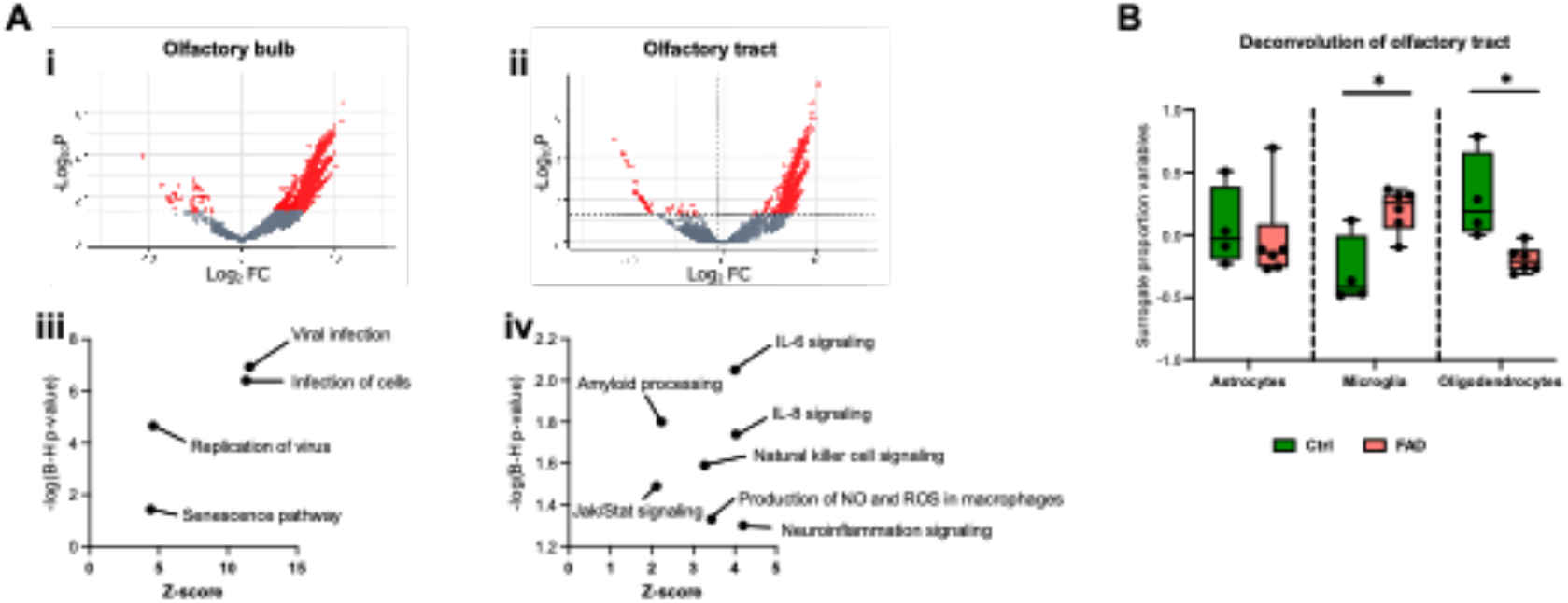
Results of TempO-Seq transcriptomic analysis in the OB and OT of FAD patients and controls. (A) Volcano plot for TempO-Seq (i, OB, ii, OT) and insight pathway analysis of transcriptional differences (iii, OB, iv, OT). (B) Deconvolution of the transcriptional differences for the OT. Asterisks are significant *p* values for post-hoc Fisher’s least significant difference.

### Spatial proteomics assay shows changes consistent with a demyelination response in the OT of FAD samples

We performed complementary nanoString spatial proteomics analysis^9^ for 88 proteins including markers for AD and immune responses (Supplemental Table 2) in tissue regions discriminated based on fluorescent markers for myelin basic protein, Aβ, Iba1 and nuclear staining (see Methods and Supplemental Table 3). The four tissue regions were high myelination and low myelination OT, and glomerular layer/external plexiform layer and granule cell layer in the OB (Figure 3A, Supplemental Figure 1). nanoString proteomics showed spatial protein specificity involved in immune and AD responses in the OT and to a lesser extent in the OB (Figure 3B). Interestingly, both the immune and AD marker responses differ between regions. For example the increase in AD markers in FAD was most marked in the low myelinated OT, and there were increases in immune markers in both low and high myelinated OT, but they were heterogeneous between these two brain regions and much lower in the OB indicating that the immune response differs between brain areas in the proximal olfactory system (Figure 3B). Strikingly, principal component analysis (PCA) of the spatial proteomics results shows orthogonal localization of FAD versus control proteomics regardless of the tissue region (Figure 3C). Furthermore, there were marked differences in protein expression between brain regions within the experimental groups (FAD and control, Figure 3D and Supplemental Figure 2). Interestingly, within the FAD cohort a large group of proteins involved in response to demyelination were expressed at high levels in high myelination OT compared to all other regions (group A in Figure 3Dii) and a group of AD markers was expressed at high levels in the low myelination OT (group B in Figure 3Dii) (eNote 1). Taken together with the deconvolution of the transcriptome in the OT showing decreased oligodendrocyte in FAD (Figure 3B) these data indicate that there is dysregulation of myelination in FAD.

**Figure 3.**
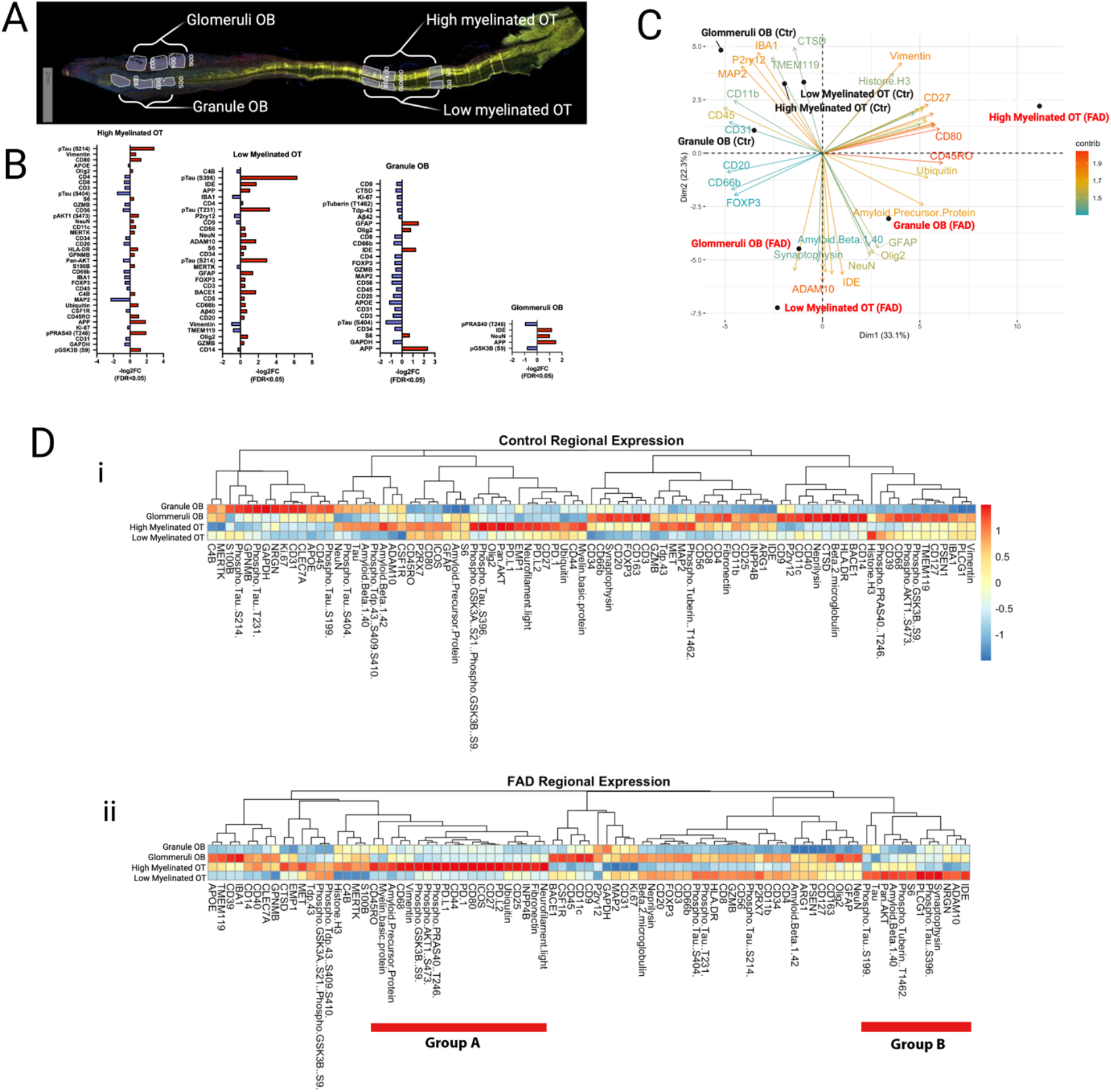
nanoString Proteomic Analysis of OB and OT Expression of Markers for AD and Immunity (A) Representative tissue section stained for myelin basic protein (MBP) showing the regions that were used for protein quantification. In the OB “Glomeruli OB” included the glomerular layer (GL) and the external plexiform layer (EPL), “Granule OB” included the granule cell layer and the accessory olfactory nucleus (AON). In the OT the high and low myelinated areas were determined by high and low MBP staining. (B) Log 2 fold change for the levels of protein expression in the different tissue regions. Red denotes higher expression in FAD. Only the proteins whose levels are significantly different between genotypes are shown. (C) Principal component analysis of the protein levels of expression. (D) Pseudocolor plots comparing between regions the levels of expression for all proteins. Proteins are segregated by Euclidean distance calculated by the total normalized reads for each protein between tissue regions or samples. i. Control tissue.ii. FAD. Group A are adjacent proteins that display higher levels in high myelinated OT and group B are adjacent proteins that display higher levels in low myelinated OT.

## Discussion

Using novel regional/spatial transcriptomic and proteomic assays, we were able to determine gene expression changes in FFPE olfactory tissues from control and FAD samples. Our findings reveal a transcriptomic signature for viral infection in the OB coupled with inflammation and dysregulation of myelination in the OT in FAD samples compared to control samples; results were confirmed by proteomic analysis. While our studies only identified human transcripts/proteins and were not designed to identify specific viruses, the most likely viruses contributing to olfactory system infection and potential CNS dysfunction are the 2 neurotropic alphaherpesviruses, herpes simplex virus type-1 (HSV-1) and varicella zoster virus (VZV). HSV-1 and VZV are strong candidates for being initiators or accelerators of AD because they increase dementia risk and elicit the same pathological characteristics of AD, including amyloid accumulation, neuroinflammation, neurodegeneration, and cognitive impairment^21-23^. Furthermore, this family of alphaherpesviruses is latent in trigeminal ganglia which innervates the OE/OB^24^, providing a direct internal route of virus entry during reactivation. Our studies find that even in patients predisposed to AD due to a genetic mutation of PS1 there is evidence of viral infection in the OB suggesting that viral infection and inflammatory response in the OT are part of the etiology of the disease. Taken together with a parallel body of literature indicating that early AD is characterized by smell loss^1-3^, amyloid deposition in the olfactory epithelium (OE), and olfactory sensory neuron (OSN) dysfunction^4, 5^, our study raises the possibility that viral infection of the OB/OT accelerates AD. Because sniff-induced beta and theta-coupled gamma oscillations generated in the OB are directionally coupled to the hippocampus^25-28^, smell loss would result in decreased hippocampal gamma oscillations that have been postulated to lead to neurodegeneration and cognitive decline^29-31^. These studies suggest that virus disruption of olfactory pathways can accelerate AD cognitive decline (Figure 4).

**Figure 4.**
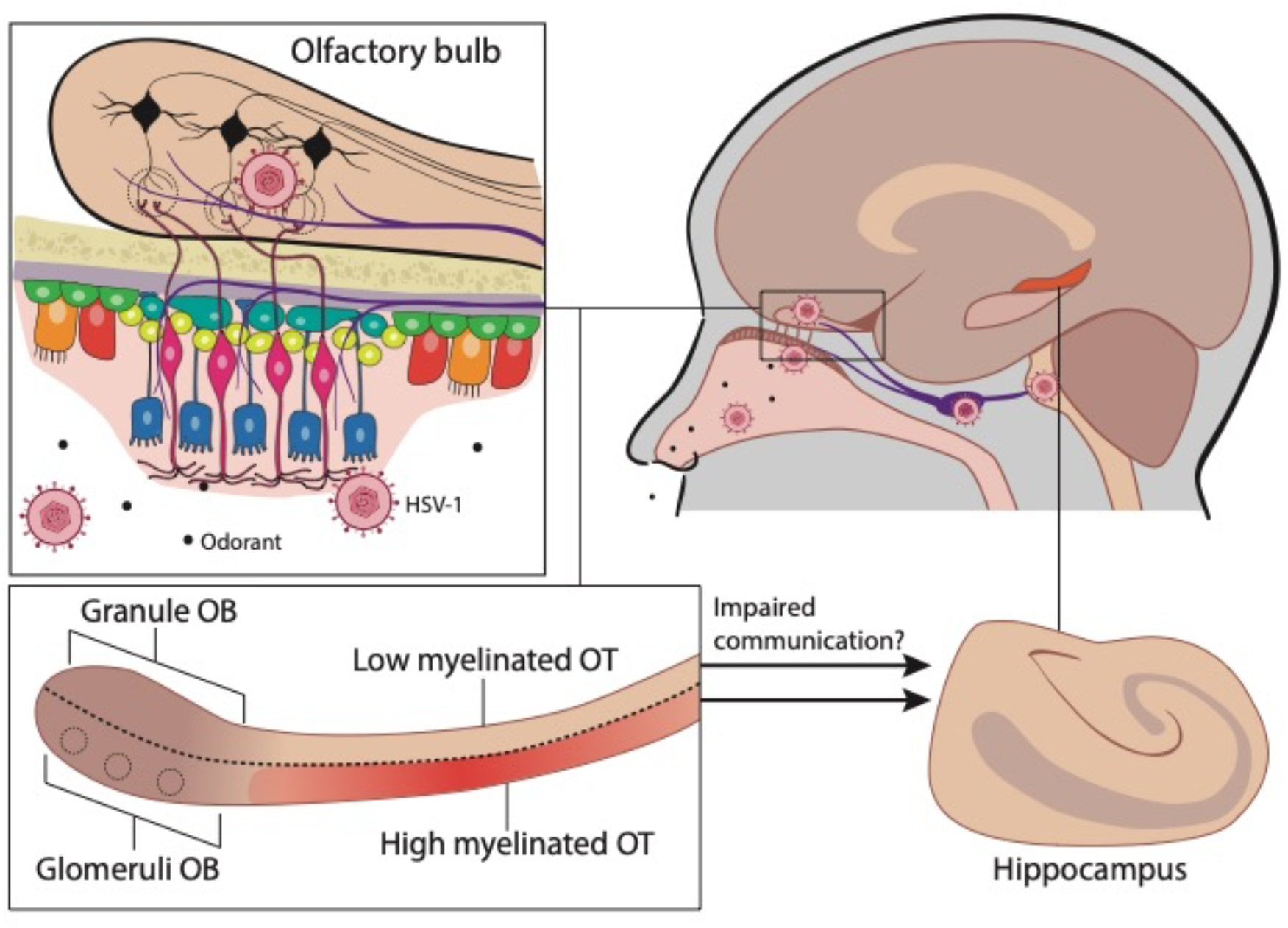
Diagram illustrating the finding of viral infection of the OB and dysregulation of myelination in the OT of FAD patients.

A limitation in our study was that we only probed for human transcripts and proteins and although strong host viral signaling pathways were identified, whether this is a broad anti-viral response or representative of a specific pathogen is unknown. Follow-up studies investigating specific viral species, such as alphaherpesviruses, in the olfactory system of AD samples are warranted. However, this may be challenging in late-stage tissues harvested at autopsies because viral infection will have likely been cleared yet chronic inflammatory processes representative of an anti-viral response may persist. Given that both HSV-1 and VZV are latent in sensory neurons of the TG in addition to the innervation of these neurons into the OE and OB provides a plausible route of recurrent infection throughout our lifespan. Because viral encephalitis is not observed at high levels in patients with AD, it is likely these pathogens are suppressed early on in the OB and OT. Overall, the transcriptional profile we have observed in this study may be representative of a routinely barraged olfactory system by pathogens and the resulting pathological ramifications such as amyloid deposition, microglia activation, and potentially myelination alterations.

## Supporting information

Supplementary Table 4 Differentially expressed genes olfactory bulb

Supplementary Table 5 Differentially expressed genes olfactory tract

Supplementary table 6 Specific pathways

## Acknowledgements

The authors thank Cathy Allen for manuscript preparation. TempO-Seq and nanoString graphics were created with BioRender.com.

## Supplemental Materials

### eNote 1. Annotations regarding the function of proteins in groups A and B of the nanoString (Fig. 3Dii)

### nanoString group A proteins: high expression in FAD high myelination OT (Fig. 3Dii)

The majority of these proteins are involved in demyelination or are axonal proteins.

#### Neurofilament light

Neurofilament light chain (NfL) is a neuronal cytoplasmic protein highly expressed in large calibre myelinated axons ^1^.

#### Fibronectin (FN1)

In toxin-induced lesions undergoing efficient remyelination, fibronectin expression was transiently increased within demyelinated areas and declined as remyelination proceeded^2^.

#### INPP4B

EAE31 is a locus controlling latency of <u>motor evoked potentials</u> (MEPs) and clinical onset of <u>experimental autoimmune encephalomyelitis</u>. By combining congenic mapping, *<u>in silico</u>* haplotype analyses, and comparative genomics Lemcke and co-workers identified <u>inositol</u> polyphosphate-4-phosphatase, type II (*Inpp4b*) as the quantitative trait gene for EAE31^3^.

#### CD25 (IL2RA)

Depletion of CD4^+^CD25^+^ cells *in vivo* facilitated the expansion of PLP reactive cells with production of T helper 1 cytokines in EAE-resistant B10.S mice. Furthermore, anti-CD25 Ab treatment before immunization resulted in EAE induction in these otherwise resistant mice. These data indicate an important role for autoantigen-specific CD4^+^CD25^+^ cells in genetic resistance to autoimmunity^4^.

#### Ubiquitin

The vast majority of cellular proteins are degraded by the 26S proteasome after their ubiquitination. Belogurov et al. ^5^ report that the major component of the myelin multilayered membrane sheath, myelin basic protein (MBP), is hydrolyzed by the 26S proteasome in a ubiquitin-independent manner both *in vitro* and in mammalian cells^5^.

#### PD.L2

Programed death ligand 2. PD-L2 is up-regulated in inflamed endothelial cells, with an intention to inhibit T-cell transmigration through the blood brain barrier^6^.

#### ICOS

The inducible costimulatory molecule (ICOS) is expressed on activated T cells and participates in a variety of important immunoregulatory functions. After the induction of experimental allergic encephalomyelitis in SJL mice with proteolipid protein (PLP), brain ICOS mRNA and protein were up-regulated on infiltrating CD3+ T cells before disease onset.^7^

#### CD80

CD4+ T Cell Expressed CD80 Regulates Central Nervous System Effector Function and Survival during Experimental Autoimmune Encephalomyelitis^8^.

#### PD.1 and PD.L1

The review by Cencioni et al describes the roles of the PD-1/ PDL-1 pathway in cancer and autoimmune diseases, especially in multiple sclerosis, and how manipulating PD-1 can be a therapeutic approach in multiple sclerosis^9^.

#### CD44

CD44 overexpression is thought to cause inflammation-independent demyelination and dysmyelination^10^.

#### Phospho-PRAS40..T246. PRAS40

(Proline-rich AKT1 substrate 1), also known as Akt1S1 and p39, is a 40-42 kDa cytoplasmic phosphoprotein that lacks generally recognized structural motifs. It is widely expressed and is considered to be key regulator of mTORC1 (mTOR plus Raptor and G beta L), a complex through which Akt signals into the cell. Through phosphorylation, mTORC1 activity is upregulated by PRAS40^11^.

#### Phospho-AKT1..S473

Involved in the dual function of the PI3K-Akt-mTORC1 axis in myelination of the peripheral nervous system^12^.

#### Phsopho-GSK3B.S9

MAI-dependent phosphorylation and inactivation of GSK3beta regulate phosphorylation of CRMP4, a cytosolic regulator of myelin inhibition, and its ability to complex with RhoA^13^.

#### Vimentin

Increase in tissue stiffness elicited by chronic demyelination of the corpus callosum is accompanied by astrogliosis, as shown by elevated GFAP and vimentin staining^14^.

#### CD68

This is a marker for macrophages. Myelin loss along with axonal destruction, the pathological hallmark of metachromatic leukodystrophy is thought to be caused by critical sulphatide levels in oligodendrocytes and Schwann cells. Immunolabelling with MBP and CD68 showed a gradient of demyelination from near-intact U-fibres to myelin-depleted white matter with diffuse macrophage infiltration.^15^.

#### APP

Data by Truong et al. identified APP and APLP2 as modulators of normal myelination and demyelination/remyelination conditions. Deletion of APP and APLP2 identifies novel interplays between the BACE1 substrates in the regulation of myelination^16^.

#### Myelin basic protein

MBP is a protein found in the myelin sheath.

#### CD45RO

Dual expression of CD45RA and CD45RO isoforms on myelin basic protein-specific CD4+ T-cell lines in multiple sclerosis^17^

### nanoString group B proteins: high expression in FAD low myelination OT Fig. 3Dii)

These proteins are AD markers.

#### IDE

Insulin-degrading-enzyme plays a crucial role in the clearance of amyloid-β and has been proposed as a therapeutical target for Alzheimer’s disease^18^.

#### ADAM10

ADAM10 is involved in the proteolytic processing of the amyloid precursor protein^19^. ADAM10 also cleaves the ectodomain of the triggering receptor expressed on myeloid cells 2 (TREM2), to produce soluble TREM2 (sTREM2), which has been proposed as a CSF and sera biomarker of neurodegeneration^20^.

#### NRGN

Neurogranin concentration in cerebrospinal fluid (CSF) is proposed as marker for synaptic dysfunction in age-related neurodegeneration^21^, and has been shown to be specifically increased in patients with Alzheimer’s disease^22^.

#### Synaptophysin

In the TMEV model, only a few large- to medium-sized synaptophysin/APP-positive bulbs were found in demyelinated areas. In MS patient tissue samples, the bulbs appeared exclusively at the inflammatory edges of lesions. In conclusion, our data suggest that synaptophysin as a reliable marker of axonal damage in the CNS in inflammatory/demyelinating conditions^23^.

#### PLCG1

Phospholipase C, gamma 1 gene mutations and abnormal splicing of PLCγ1 gene has been identified in AD using both high-throughput screening data and a deep learning-based prediction^24^.

#### S6

Ribosomal protein S6

#### Phospho Tuberin T1462

The control of translation is disturbed in Alzheimer’s disease (AD). Morel et al.^25^ analyzed the crosslink between the up regulation of double-stranded RNA-dependent-protein kinase (PKR) and the down regulation of mammalian target of rapamycin (mTOR) signaling pathways via p53, the protein Regulated in the Development and DNA damage response 1 (Redd1) and the tuberous sclerosis complex (TSC2) factors in two beta-amyloid peptide (Abeta) neurotoxicity models. In SH-SY5Y cells, Abeta42 induced an increase of P(T451)-PKR and of the ratio p66/(p66+p53) in nuclei and a physical interaction between these proteins. Redd1 gene levels increased and P(T1462)-TSC2 decreased. These disturbances were earlier in rat primary neurons with nuclear co-localization of Redd1 and PKR. The PKR gene silencing in SH-SY5Y cells prevented these alterations. p53, Redd1 and TSC2 could represent the molecular links between PKR and mTOR in Abeta neurotoxicity. PKR could be a critical target in a therapeutic program of AD.

#### Tau, Phospho.Tau..S199

Phosphorylated tau.

---

**Supplementary Table 1.** Antibody Information for Multispectral Immunohistochemistry

| Target | Concentration | Antigen retrieval conditions | Vendor | Catalogue number | Host Species |
| --- | --- | --- | --- | --- | --- |
| Ab | 1:500 | EDTA ER2 pH9 | Biolegend | 803001 | Mouse |
| Iba1 | 1:10,000 | EDTA ER2 pH9 | Wako | 019-19741 | Rabbit |
| GFAP | 1:500 | EDTA ER2 pH9 | Dako/Agilent | Z033429-2 | Rabbit |
| Cleaved Caspase 3 | 1:100 | EDTA ER2 pH9 | Cell Signaling | 9664L | Rabbit |
| p-Tau | 1:200 | Citrate ER1 pH6 | Invitrogen | MN1020 | Mouse |
| DCX | 1:5,000 | Citrate ER1 pH6 | Abcam | ab18723 | Rabbit |
| CD68 | 1:500 | Citrate ER1 pH6 | Dako/Agilent | M0814 | Mouse |
| PLP | 1:400 | Citrate AR buffer + TritonX100 pH6 | PLP antibody source was<sup>10</sup> |  | Rat |

Antigen retrieval was performed on a fully automated immunostainer using either ER1 (citrate with a pH range of 5.9-6.1) or ER2 (EDTA with a pH range of 8.9-9.1) buffers, except for PLP which was manually processed in the microwave with citrate AR buffer and TritonX100 (pH6).

**Supplementary Table 2.** Antibodies Used for nanoString

| Item name | Item Number |
| --- | --- |
| GeoMx Human Protein Core for NGS | 121300129 |
| GeoMx Immune Cell Typing Assay | 121300130 |
| GeoMx Immune Activation Status Assay | 121300132 |
| GeoMx Myeloid Assay | 121300137 |
| GeoMx PI3K/AKT Signaling Assay | 121300136 |
| GeoMx Neural Cell Typing Assay | 121300138 |
| GeoMx Alzheimer's Pathology Assay | 121300139 |
| GeoMx Glial Cell Subtyping Assay | 121300143 |

**Supplementary Table 3.** Antibody Information for Immunofluorescence for nanoString

| Antibody ID | Name | company | clone # | Catalog # | concentration used |
| --- | --- | --- | --- | --- | --- |
| syto13 | syto13 | NanoString |  | 121300306 | 25 |
| MF-304 | Beta amyloid | Novus | MOAB-2 | NBP2-13075AF532 | 1:100 |
| MF-618 | MBP | Novus | 2H9 | NBP2-22121AF594 | 1:100 |
| MF-643 | Iba1 | millipore | 20A12.1 | MABN92-AF647 | 1:100 |

**Supplementary Table 4.** Excel worksheet with TempO-Seq raw counts and p-values for all genes for the olfactory bulb (control vs. FAD)

**Supplementary Table 5.** Excel worksheet with TempO-Seq raw counts and p-values for all genes for the olfactory tract (control vs. FAD)

**Supplementary Table 6.** Excel worksheet with genes included in specific pathways by Insight analysis.

**Supplementary Figure 1.**
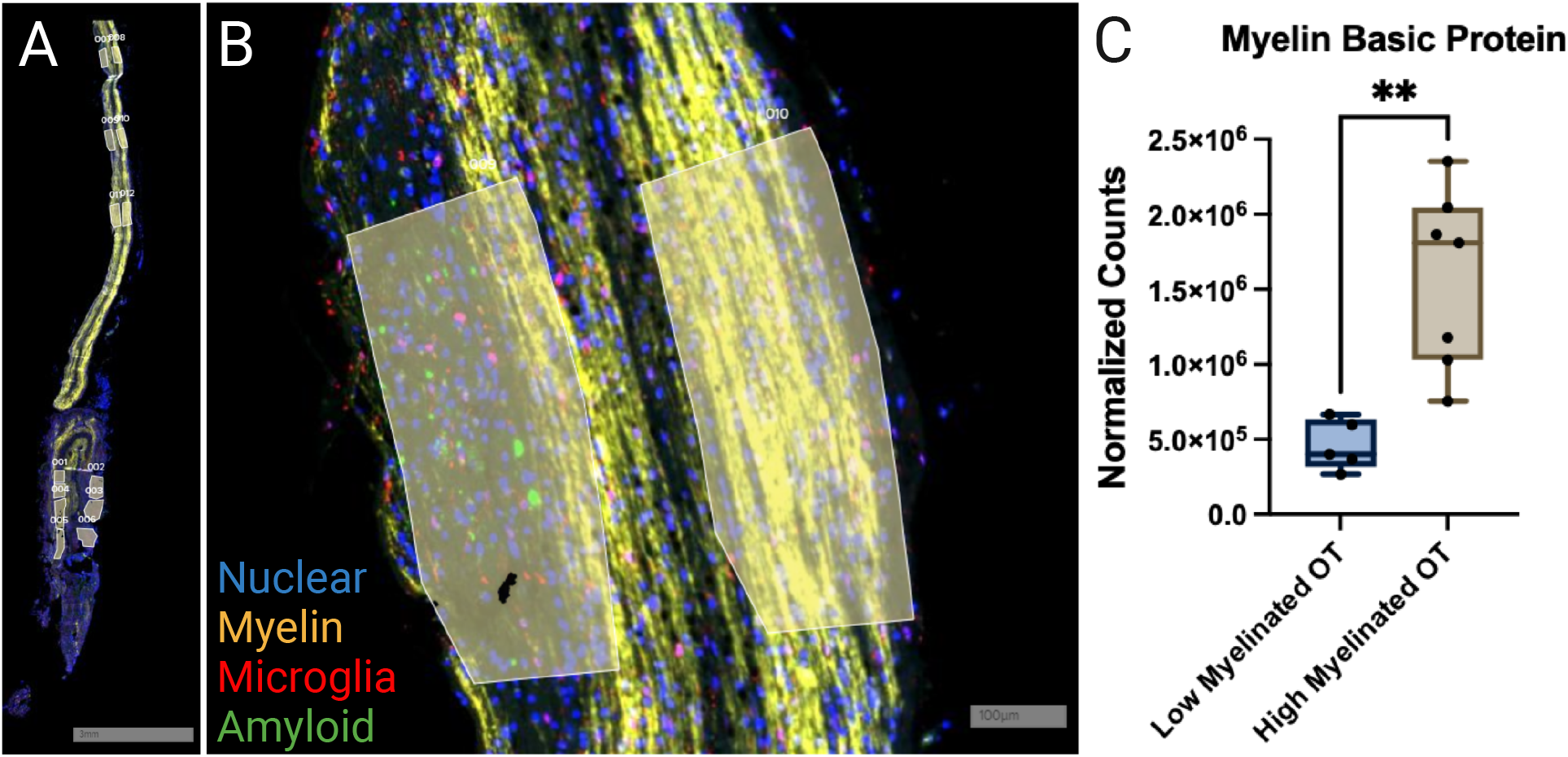
Quantification of expression of myelin basic protein in the high and low myelinated regions of the OT of FAD subjects using nanoString spatial profiling. (A) A representative horizontal tissue section (FAD 359) of the OB/OT showing regions used for protein quantification. The sections are immunolabeled for myelin basic protein, Iba1, Aβ and nuclear staining (styo13). (B) Closeup of OT areas with high and low myelination. (C) Normalized counts for myelin basic protein in the low and high myelination areas. The difference is statistically significant (p<0.05, pairwise t-test).

**Supplementary Figure 2.**
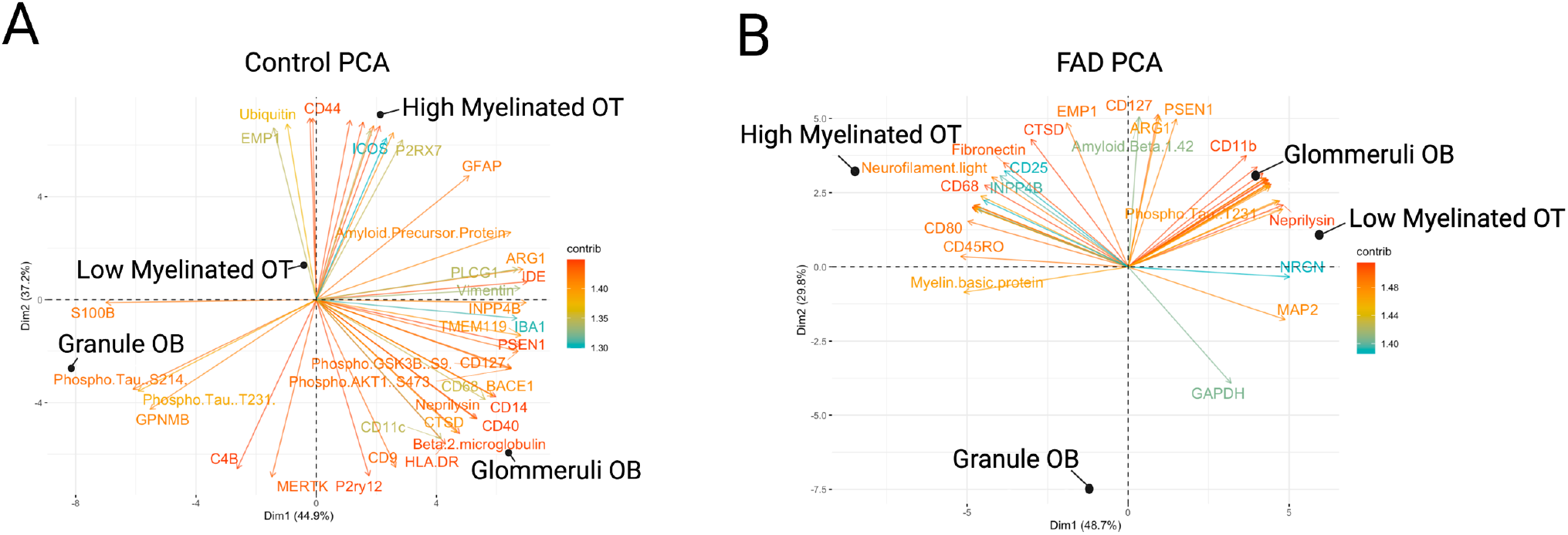
Principal component analysis of the proteomic data for the control and FAD datasets. A *principal component analysis of nanoString* protein expression shows increased distance between high and low myelinated OT in the FAD tissue. (A) Control. (B) FAD.

## References

1. Wheeler PL, Murphy C. Olfactory Measures as Predictors of Conversion to Mild Cognitive Impairment and Alzheimer’s Disease. Brain Sciences 2021;11.

2. Murphy C. Olfactory and other sensory impairments in Alzheimer disease. Nat Rev Neurol 2019;15:11–24.

3. Dibattista M, Pifferi S, Menini A, Reisert J. Alzheimer’s Disease: What Can We Learn From the Peripheral Olfactory Systemã Frontiers in Neuroscience 2020;14:440.

4. Tremblay C, Serrano GE, Intorcia AJ, et al. Olfactory Bulb Amyloid-β Correlates With Brain Thal Amyloid Phase and Severity of Cognitive Impairment. Journal of Neuropathology & Experimental Neurology 2022;81:643–649.

5. Wesson DW, Levy E, Nixon RA, Wilson DA. Olfactory Dysfunction Correlates with Amyloid-β Burden in an Alzheimer’s Disease Mouse Model. The Journal of Neuroscience 2010;30:505.

6. Attems J, Walker L, Jellinger KA. Olfactory bulb involvement in neurodegenerative diseases. Acta Neuropathologica 2014;127:459–475.

7. Acosta-Baena N, Sepulveda-Falla D, Lopera-Gómez CM, et al. Pre-dementia clinical stages in presenilin 1 E280A familial early-onset Alzheimer’s disease: a retrospective cohort study. The Lancet Neurology 2011;10:213–220.

8. Trejo CL, Babic M, Imler E, et al. Extraction-free whole transcriptome gene expression analysis of FFPE sections and histology-directed subareas of tissue. PLoS One 2019;14:e0212031.

9. Merritt CR, Ong GT, Church SE, et al. Multiplex digital spatial profiling of proteins and RNA in fixed tissue. Nature Biotechnology 2020;38:586–599.

10. Quiroz YT, Sperling RA, Norton DJ, et al. Association Between Amyloid and Tau Accumulation in Young Adults With Autosomal Dominant Alzheimer Disease. JAMA Neurology 2018;75:548–556.

11. Parra ER, Uraoka N, Jiang M, et al. Validation of multiplex immunofluorescence panels using multispectral microscopy for immune-profiling of formalin-fixed and paraffin-embedded human tumor tissues. Scientific Reports 2017;7:13380.

12. Bubak AN, Como CN, Hassell JE, et al. Targeted RNA Sequencing of VZV-Infected Brain Vascular Adventitial Fibroblasts Indicates That Amyloid May Be Involved in VZV Vasculopathy. Neurology - Neuroimmunology Neuroinflammation 2022;9:e1103.

13. Varet H, Brillet-Gueguen L, Coppee JY, Dillies MA. SARTools: A DESeq2- and EdgeR-Based R Pipeline for Comprehensive Differential Analysis of RNA-Seq Data. PLoS One 2016;11:e0157022.

14. Robinson MD, McCarthy DJ, Smyth GK. edgeR: a Bioconductor package for differential expression analysis of digital gene expression data. Bioinformatics 2010;26:139–140.

15. Love MI, Huber W, Anders S. Moderated estimation of fold change and dispersion for RNA-seq data with DESeq2. Genome Biol 2014;15:550.

16. Yu G, Wang L-G, Han Y, He Q-Y. clusterProfiler: an R Package for Comparing Biological Themes Among Gene Clusters. OMICS: A Journal of Integrative Biology 2012;16:284–287.

17. Yu G, Wang L-G, Yan G-R, He Q-Y. DOSE: an R/Bioconductor package for disease ontology semantic and enrichment analysis. Bioinformatics 2015;31:608–609.

18. Yamamura T, Konola JT, Wekerle H, Lees MB. Monoclonal Antibodies Against Myelin Proteolipid Protein: Identification and Characterization of Two Major Determinants. Journal of Neurochemistry 1991;57:1671–1680.

19. Kohl Z, Schlachetzki JCM, Feldewerth J, et al. Distinct Pattern of Microgliosis in the Olfactory Bulb of Neurodegenerative Proteinopathies. Neural Plasticity 2017;2017:3851262.

20. Condello C, Yuan P, Schain A, Grutzendler J. Microglia constitute a barrier that prevents neurotoxic protofibrillar Aβ42 hotspots around plaques. Nature Communications 2015;6:6176.

21. Eimer WA, Vijaya Kumar DK, Navalpur Shanmugam NK, et al. Alzheimer’s Disease-Associated beta-Amyloid Is Rapidly Seeded by Herpesviridae to Protect against Brain Infection. Neuron 2018;99:56–63 e53.

22. Readhead B, Haure-Mirande JV, Funk CC, et al. Multiscale Analysis of Independent Alzheimer’s Cohorts Finds Disruption of Molecular, Genetic, and Clinical Networks by Human Herpesvirus. Neuron 2018;99:64–82 e67.

23. Itzhaki RF, Lathe R. Herpes Viruses and Senile Dementia: First Population Evidence for a Causal Link. J Alzheimers Dis 2018;64:363–366.

24. Schaefer ML, Bottger B, Silver WL, Finger TE. Trigeminal collaterals in the nasal epithelium and olfactory bulb: a potential route for direct modulation of olfactory information by trigeminal stimuli. J Comp Neurol 2002;444:221–226.

25. Martin C, Beshel J, Kay LM. An olfacto-hippocampal network is dynamically involved in odor-discrimination learning. JNeurophysiol 2007;98:2196–2205.

26. Gourevitch B, Kay LM, Martin C. Directional coupling from the olfactory bulb to the hippocampus during a go/no-go odor discrimination task. J Neurophysiol 2010;103:2633–2641.

27. Nguyen Chi V, Muller C, Wolfenstetter T, et al. Hippocampal Respiration-Driven Rhythm Distinct from Theta Oscillations in Awake Mice. J Neurosci 2016;36:162–177.

28. Pena RR, Medeiros DdC, Guarnieri LdO, et al. Home-cage odors spatial cues elicit theta phase/gamma amplitude coupling between olfactory bulb and dorsal hippocampus. Neuroscience 2017;363:97–106.

29. Iaccarino HF, Singer AC, Martorell AJ, et al. Gamma frequency entrainment attenuates amyloid load and modifies microglia. Nature 2016;540:230–235.

30. Gillespie Anna K, Jones Emily A, Lin Y-H, et al. Apolipoprotein E4 Causes Age-Dependent Disruption of Slow Gamma Oscillations during Hippocampal Sharp-Wave Ripples. Neuron 2016;90:740–751.

31. Salimi M, Tabasi F, Abdolsamadi M, et al. Disrupted connectivity in the olfactory bulb-entorhinal cortex-dorsal hippocampus circuit is associated with recognition memory deficit in Alzheimer’s disease model. Scientific Reports 2022;12:4394.

## References for eNote 1

1. Gaetani, L., et al. Neurofilament light chain as a biomarker in neurological disorders. Journal of Neurology, Neurosurgery &amp;amp; Psychiatry 90, 870 (2019).

2. Stoffels, J.M.J., et al. Fibronectin aggregation in multiple sclerosis lesions impairs remyelination. Brain 136, 116–131 (2013).

3. Lemcke, S., et al. Nerve Conduction Velocity Is Regulated by the Inositol Polyphosphate-4-Phosphatase II Gene. The American Journal of Pathology 184, 2420–2429 (2014).

4. Reddy, J., et al. Myelin proteolipid protein-specific CD4&lt;sup&gt;+&lt;/sup&gt;CD25&lt;sup&gt;+&lt;/sup&gt; regulatory cells mediate genetic resistance to experimental autoimmune encephalomyelitis. Proceedings of the National Academy of Sciences of the United States of America 101, 15434 (2004).

5. Belogurov, A., Jr., et al. Multiple Sclerosis Autoantigen Myelin Basic Protein Escapes Control by Ubiquitination during Proteasomal Degradation *. Journal of Biological Chemistry 289, 17758–17766 (2014).

6. Zhao, S., Li, F., Leak, R.K., Chen, J. & Hu, X. Regulation of Neuroinflammation through Programed Death-1/Programed Death Ligand Signaling in Neurological Disorders. Frontiers in Cellular Neuroscience 8(2014).

7. Rottman, J.B., et al. The costimulatory molecule ICOS plays an important role in the immunopathogenesis of EAE. Nature Immunology 2, 605–611 (2001).

8. Podojil, J.R., Kohm, A.P. & Miller, S.D. CD4&lt;sup&gt;+&lt;/sup&gt; T Cell Expressed CD80 Regulates Central Nervous System Effector Function and Survival during Experimental Autoimmune Encephalomyelitis. The Journal of Immunology 177, 2948 (2006).

9. Cencioni, M.T. The immune regulation of PD-1/PDL-1 axis, a potential biomarker in multiple sclerosis. Neuroimmunology and Neuroinflammation 7, 277–290 (2020).

10. Tuohy, T.M., et al. CD44 overexpression by oligodendrocytes: a novel mouse model of inflammation-independent demyelination and dysmyelination. Glia 47, 335–345 (2004).

11. Bercury, K.K., et al. Conditional ablation of raptor or rictor has differential impact on oligodendrocyte differentiation and CNS myelination. J Neurosci 34, 4466–4480 (2014).

12. Figlia, G., Norrmén, C., Pereira, J.A., Gerber, D. & Suter, U. Dual function of the PI3K-Akt-mTORC1 axis in myelination of the peripheral nervous system. eLife 6, e29241 (2017).

13. Alabed, Y.Z., Pool, M., Ong Tone, S., Sutherland, C. & Fournier, A.E. GSK3β Regulates Myelin-Dependent Axon Outgrowth Inhibition through CRMP4. The Journal of Neuroscience 30, 5635 (2010).

14. Urbanski, M.M., Brendel, M.B. & Melendez-Vasquez, C.V. Acute and chronic demyelinated CNS lesions exhibit opposite elastic properties. Scientific Reports 9, 999 (2019).

15. Ponath, G., et al. Myelin phagocytosis by astrocytes after myelin damage promotes lesion pathology. Brain 140, 399–413 (2017).

16. Truong, P.H., et al. Amyloid precursor protein and amyloid precursor-like protein 2 have distinct roles in modulating myelination, demyelination, and remyelination of axons. Glia 67, 525–538 (2019).

17. Qin, Y., van den Noort, S., Kurt, J. & Gupta, S. Dual expression of CD45RA and CD45RO isoforms on myelin basic protein-specific CD4+ T-cell lines in multiple sclerosis. Journal of Clinical Immunology 13, 152–161 (1993).

18. Kurochkin, I.V., Guarnera, E. & Berezovsky, I.N. Insulin-Degrading Enzyme in the Fight against Alzheimer’s Disease. Trends in Pharmacological Sciences 39, 49–58 (2018).

19. Haass, C., Kaether, C., Thinakaran, G. & Sisodia, S. Trafficking and proteolytic processing of APP. Cold Spring Harb Perspect Med 2, a006270 (2012).

20. Yang, J., et al. TREM2 ectodomain and its soluble form in Alzheimer’s disease. Journal of Neuroinflammation 17, 204 (2020).

21. Casaletto, K.B., et al. Neurogranin, a synaptic protein, is associated with memory independent of Alzheimer biomarkers. Neurology 89, 1782 (2017).

22. Willemse, E.A.J., et al. Neurogranin as Cerebrospinal Fluid Biomarker for Alzheimer Disease: An Assay Comparison Study. Clinical Chemistry 64, 927–937 (2018).

23. Gudi, V., et al. Synaptophysin Is a Reliable Marker for Axonal Damage. Journal of Neuropathology & Experimental Neurology 76, 109–125 (2017).

24. Kim, S.-H., et al. Prediction of Alzheimer’s disease-specific phospholipase c gamma-1 SNV by deep learning-based approach for high-throughput screening. Proceedings of the National Academy of Sciences 118, e2011250118 (2021).

25. Morel, M., et al. Evidence of molecular links between PKR and mTOR signalling pathways in Aβ neurotoxicity: Role of p53, Redd1 and TSC2. Neurobiology of Disease 36, 151–161 (2009).

